# Suite2p: beyond 10,000 neurons with standard two-photon microscopy

**DOI:** 10.1101/061507

**Authors:** Marius Pachitariu, Carsen Stringer, Mario Dipoppa, Sylvia Schröder, L. Federico Rossi, Henry Dalgleish, Matteo Carandini, Kenneth D. Harris

## Abstract

Two-photon microscopy of calcium-dependent sensors has enabled unprecedented recordings from vast populations of neurons. While the sensors and microscopes have matured over several generations of development, computational methods to process the resulting movies remain inefficient and can give results that are hard to interpret. Here we introduce Suite2p: a fast, accurate and complete pipeline that registers raw movies, detects active cells, extracts their calcium traces and infers their spike times. Suite2p runs on standard workstations, operates faster than real time, and recovers ~2 times more cells than the previous state-of-the-art method. Its low computational load allows routine detection of ~10,000 cells simultaneously with standard two-photon resonant-scanning microscopes. Recordings at this scale promise to reveal the fine structure of activity in large populations of neurons or large populations of subcellular structures such as synaptic boutons.

## Introduction

Standard resonance-scanning two-photon microscopes readily image the activity of large numbers of neurons, but pipelines for processing the resulting data still suffer from significant limitations. Ideally, such a pipeline should satisfy several criteria. First, it should be fast, to keep up with ever-larger data sets produced by next-generation microscopes ^1,2^. Second, the pipeline should be transparent and its results interpretable, so that the original data undergo minimal processing, and a human curator can recognize mistakes or biases in the pipeline’s output. Third, the pipeline should be accurate, so that its results require only brief curation by a human operator. Fourth, the pipeline should generalize to recordings of multiple cell types, subcellular structures, and brain regions, which can exhibit widely different activity patterns. Fifth, the pipeline should appropriately model and handle experimental confounds such as neuropil contamination ^3^. Finally, it would be ideal if the pipeline could run on inexpensive workstations rather than requiring a cluster of servers, as some current software packages do ^4^.

To fulfil these criteria, we developed Suite2p, an end-to-end pipeline made of fast and accurate algorithms (Figure 1a). The pipeline involves four independent stages: 1) image registration; 2) region-of-interest (ROI) detection; 3) ROI labelling and quality control; 4) activity extraction with neuropil correction and spike deconvolution (Figure 1b-e). Each stage can be separately adapted to new types of data as needs arise, unlike approaches that combine multiple steps into a single, less transparent procedure.

**Figure 1.**
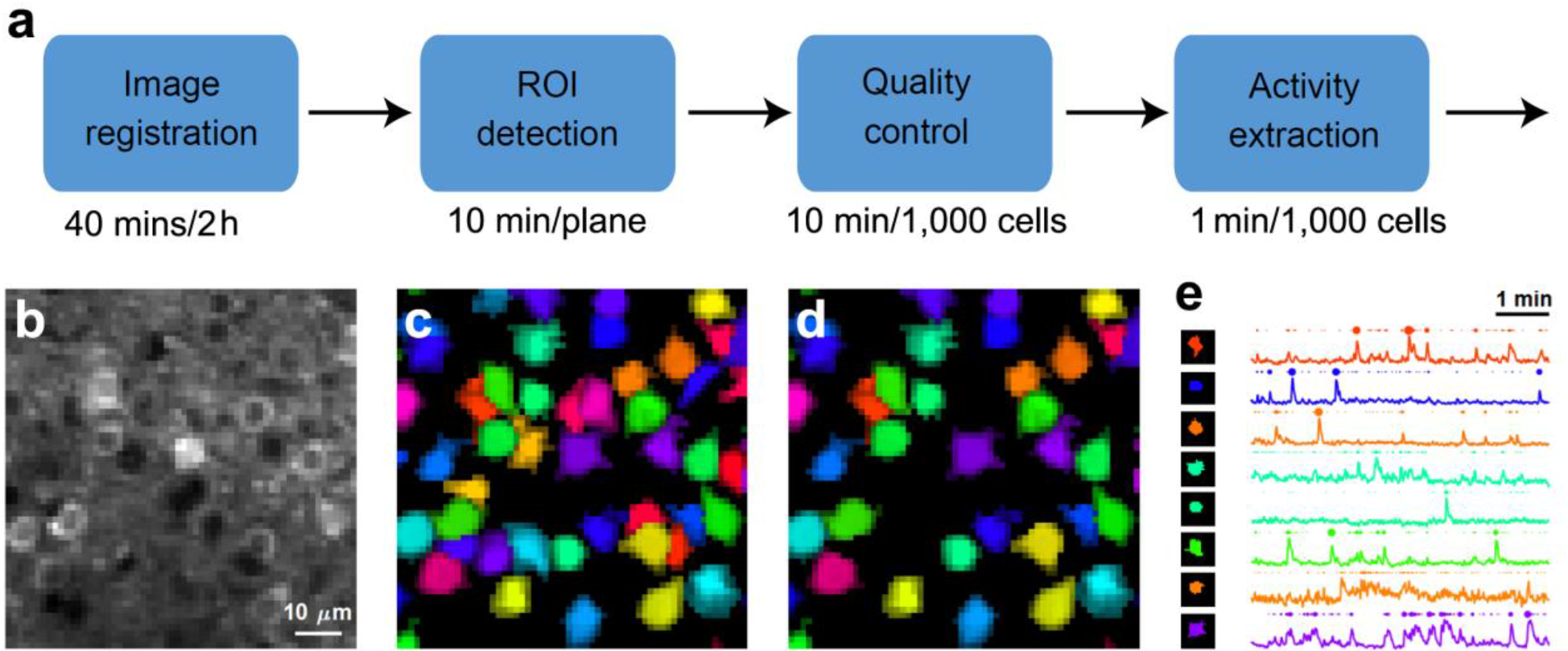
Pipeline schematic. (a) The Suite2p pipeline involves four steps: image registration, ROI detection, quality control (i.e. detection of ROIs corresponding to somata), and activity detection (removal of neuropil contamination plus optional spike deconvolution). The processing time required for each step is indicated, based on a 2 hour recording of 9623 neurons, 10 planes over a field of view of 900μm x 900μm, where each plane is sampled at 3 Hz. (b) Mean fluorescence image of a small portion of the field of view after sub-sample image registration. (c) Results of the ROI detection stage for this portion of the field of view. Colors indicate cluster assignments of pixels assigned to ROIs (chosen randomly for each ROI). (d) Quality control by manual curation and removal of non-somatic ROIs (white). (e) Neuropil-corrected activity traces for a subset of the cells in d, together with their deconvolved spike times, represented as dots of area proportional to firing rate.

Suite2p requires substantially less computational power than previous approaches ^5–9^ allowing it to provide state-of-the-art cell yields and accuracy faster than real-time on a $1,000 workstation. The software package is available at github.com/cortex-lab/Suite2P. In addition to the data processing pipeline, the package includes a user-friendly graphical user interface (GUI) for manual inspection and curation.

As well as faster running speed, Suite2p offers substantial computational advantages over existing algorithms. For instance, unlike algorithms based on independent component analysis ^7^, it does not assume that a cell’s activity should be independent of the surrounding neuropil (while in fact they can be strongly correlated). Moreover, Suite2p finds about twice as many neurons as a widely used method^5^ based on Constrained Non-negative Matrix Factorization (CNMF). Suite2p may also be preferable to methods that rely on a single anatomical image ^10,11^, as these methods tend to miss cells that are sparsely active and have low baseline fluorescence (a common occurrence in the cerebral cortex). Finally, Suite2p appears to be preferable to algorithms based on greedily segmenting neighboring pixels of high correlation ^1,8,12,13^ as such methods miss many cells.

## Results

The Suite2p pipeline comprises four stages, each of which has clearly defined inputs and interpretable outputs, and can be run independently (Figure 1a). Stage 1 is image registration: it takes as input the raw calcium movie, and outputs a registered movie (Figure 1b). Stage 2 is ROI detection: it takes as input the registered movie and outputs a set of spatial ROIs with positive weights for each pixel (Figure 1c). Stage 3 is quality control: it takes as inputs the ROI shapes and several statistics computed from the activity traces, and labels each ROI as a cell or not a cell (Figure 1d). Users can manually over-ride these labels in a GUI, which provides training data to the automated classifier we use to assign these labels. Stage 4 is activity and neuropil extraction, with optional spike deconvolution (Figure 1e).

For each stage, we first describe and address the challenges arising from constraints of the recording method, and then validate the pipeline using a combination of benchmarking strategies. Finally, we illustrate the performance of the pipeline by applying it to standard two-photon data acquired in the primary visual cortex of the awake mouse. With these datasets, we show that Suite2p can rapidly and precisely reveal the activity of >10,000 neurons and of large populations of dendritic spines and axonal boutons.

### Stage 1: image registration via phase correlation

The first stage involves correcting for the effects of brain movement, by registering all frames in the movie to each other.

Common registration techniques used in two-photon microscopy rely on finding the cross-correlation peak between a frame and a target image ^14^. This peak can be computed efficiently with the fast Fourier transform (FFT) and determines the XY offset (not necessarily an integer) by which the frame should be shifted. The frame is then shifted using FFT-based interpolation.

A disadvantage of this approach is that it is driven by the low spatial frequencies that dominate images, to the expense of the high-frequency content that is essential for registration. At typical magnifications, this implies ignoring somata and other calcium-filled cellular compartments. To emphasize the high-frequency content, we used phase correlation, which applies spatial whitening to the images before computing the cross-correlation map^15,16^. We extended this method to detect sub-pixel shifts (down to 1/10 of a pixel), by interpolating the phase correlation map near its maximum using a squared-exponential kernel (kriging).

We tested the resulting algorithm on simulated data with known translation and realistic noise (Figure 2abc) and found that it outperformed the standard method of cross-correlation (Figure 2de). For 512×512 pixel frames, our method was about as fast as standard, non-subpixel cross-correlation, and >15 times faster than 10x upsampled cross-correlation (Figure 2f). Over a wide range of simulation parameters, our method performed consistently well, with errors of ~0.1 pixels, while the standard methods lagged well behind (Figure 2g-n). Moreover, our algorithm could be further accelerated ~4 times using GPU-based computations.

**Figure 2.**
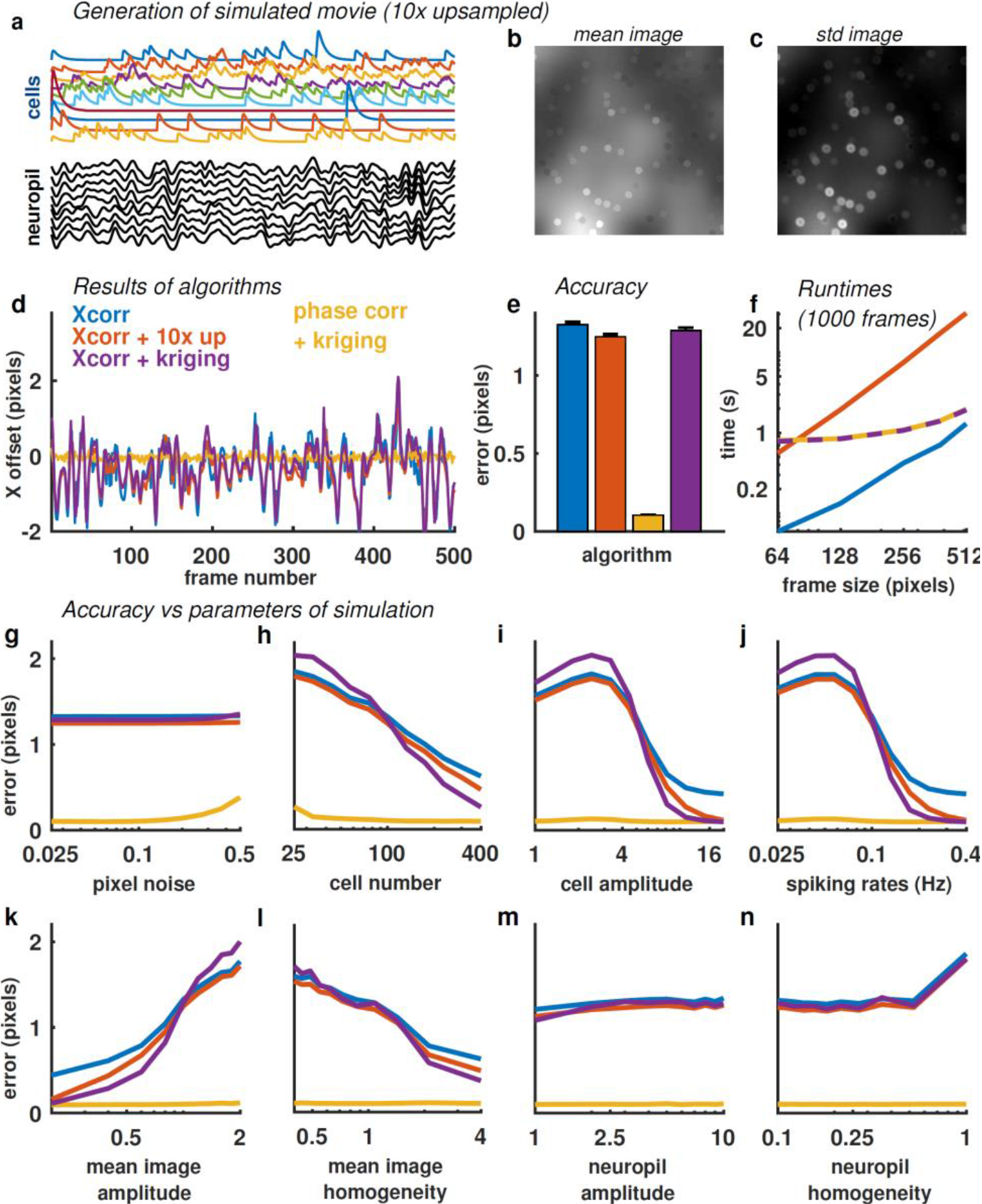
Correcting rigid motion with subpixel phase correlation. (a) Example frame from a simulation of imaging data, subject to random rigid movement and activity-dependent changes in cellular and extracellular fluorescence. The cell masks were disks distributed randomly over the field of view. (b,c) Mean and standard deviation of fluorescence signal (without image registration). (d) Residual registration error (x offset) as a function of time, for four registration algorithms (standard cross-correlation, cross-correlation with Fourier upsampling, cross-correlation with kriging interpolation and phase correlation with kriging interpolation). Phase correlation with kriging provides best performance. (e) Mean residual error for these four algorithms (averaged over 10 randomized simulations). (f) Runtimes of registration for the four algorithms. Color codes as in (d), dashed line indicates identical runtime. (g-n) Mean residual error as a function of eight simulation parameters: (g) pixel noise, (h) cell number, (i) amplitude of cell responses, (j) cell spiking rates, (k) amplitude of mean image, (l) spatial homogeneity of the mean image, (m) amplitude of the neuropil activity, and (n) homogeneity of the neuropil activity. Phase correlation with kriging provides better performance over a wide range of parameter values.

In most cases, the rigid alignment algorithm appears sufficient (movie 1), but in others there appears to be residual non-rigid motion (movie 2, left panel). Suite2p encourages users to visually inspect the output of the algorithm after rigid alignment has run, to determine whether to use an option to correct for rotational and non-rigid brain movements^17^. To compute non-rigid alignments, Suit2p divides the image in blocks, and uses phase correlation to estimate XY offsets for each block. It then interpolates these XY offsets using Gaussian basis functions centered on the midpoints of each block, thus generating a globally non-rigid transformation. The interpolated pixel shifts are then discretized and applied to the original image at the pixel level.

We have applied this non-rigid method successfully in experiments where movements were large relative to the size of the regions of interest such as cells (movie 2), boutons or spines. In our experience, the main source of apparent non-rigid motion is the non-simultaneous acquisition of lines at different Y offsets in each frame^17^. We therefore simulated movies with these properties, and validated the algorithm on those movies (movie 3, Supplementary Figure 1).

### Stage 2: ROI detection

The second stage is central to our pipeline: it takes as input the registered movie and outputs a set of spatial regions of interest (ROIs), with positive weights for each pixel (Figure 1c).

At the core of this stage is a novel source-extraction algorithm, derived from a simple model of the underlying biological structure, and of the optics of the microscope. This model assumes that the recorded signal in each pixel is a sum of signals originating from distinct active compartments, such as somata, dendrites, spines or boutons (our regions of interest, depending on magnification), and a more diffuse contamination signal originating in the neuropil.

The neuropil produces a large and diffuse contamination signal, arising from the averaged activity of a large number of out-of-focus dendrites and axons. Two-photon microscopes acquire signals with a point-spread function that is typically one order of magnitude wider in the Z-axis than in X and Y. Thus, the signal recorded in a single plane is a weighted average of a volume extending across tens of microns in Z. The fluorescence originating in this volume arises mainly from neuropil (i.e. axons and dendrites), and it contaminates the signals measured from individual cells, even if those cells might appear to be the only element occupying a spatial position ^3^. It is essential for a modeling framework to account for this contamination, because it contributes a large amount of variance to the overall recorded signals, and is far from independent from the activity of the desired regions of interest ^20^.

To account for the neuropil signal, we must first understand its spatial distribution. By analyzing pixels outside of ROIs measured in mouse visual cortex, we found that the neuropil signal is highly correlated to the total signal recorded at the soma of each cell, over long distances (Figure 3a, b). Cellular fluorescence transients constitute deviations from an otherwise linear relationship between a somatic signal and the surrounding neuropil (Figure 3c, d). These deviations occur primarily when the neuropil signal is itself large, reflecting increasing cell firing probability at times of high network activity. This result emphasizes how a spatially smooth neuropil signal is indeed present at the soma and accounts for a considerable fraction of the recorded signal ^3,20,21^.

**Figure 3.**
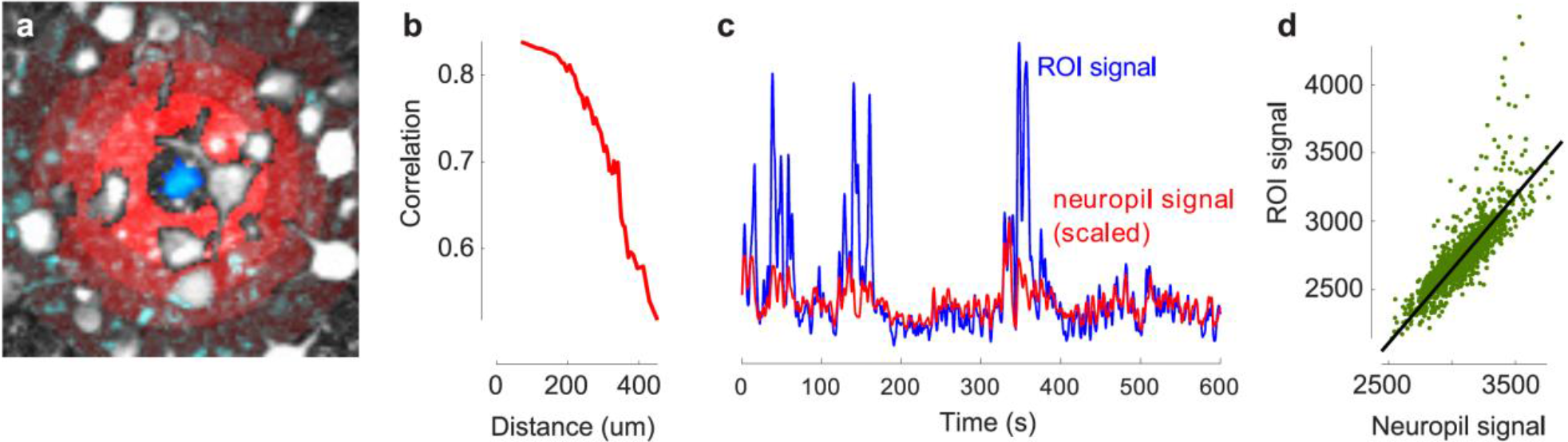
Spatial statistics of the neuropil signal. (a) Example cell (blue) and neuropil areas (red-shaded annuli) superimposed on the variance image (white), for an imaging session in superficial V1. The neuropil signal is defined as the signal inside the red-shaded annuli but not in any detected ROIs. (b) Correlation between the cell’s fluorescence signal and the neuropil signal is large, and decays slowly with distance. (c) Example timecourse of ROI signal and neuropil signal. The neuropil signal has been scaled by 0.93, as found by our algorithm. (d) Scatter plot of cell and neuropil signals (green points), together with a robust regression fit (black line), that ignores outliers (above fit line, which correspond to spike-related transients).

Guided by these results, we represented the neuropil signal in a set of spatially-localized basis functions, which allow the signal to vary slowly across space. This approach differs from previous methods, which do not explicitly model the neuropil ^7^ or model it as a one-dimensional signal shared by all pixels with different weights ^5^.

We consider a set of spatial basis functions ***B***, which together tile the full field of view, covering each pixel *k*. The neuropil signal in each basis function *j* is a smooth timecourse ***n**_j_(t)*. In the model, therefore, the neuropil signal at pixel *k* is *∑_j_**B**_kj_**n**_j_(t*). We chose the basis functions to be isotropic 2d raised cosines with a default spacing of three times the diameter of a cell.

We assumed that signals from the ROIs (somata, dendrites, spines or boutons, depending on the scale of interest) are spatially localized, and that each pixel in ROI *i* follows the same timecourse ***f**_i_(t)*, scaled by a constant factor according to a (very sparse) matrix:

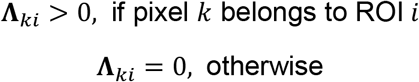

The full model for the recorded signal ***r**_k_* at pixel *k* is thus

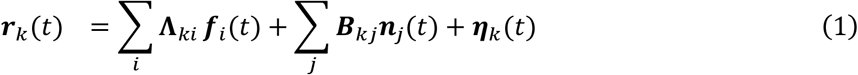

where the first term models the activity of the ROIs, the second term models the neuropil, and the third term is measurement noise, which we take to be Gaussian:

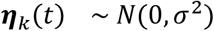

To fit the model to the data, we alternate between three steps, described in more detail in Methods:

- ROI detection, to find new sources that are not yet accounted for.
- Activity extraction, to re-estimate the timecourses of these components and the contribution of the neuropil.
- Pixel re-assignments, to re-estimate the spatial distribution of the ROIs in the context of all other ROIs.

We typically run the procedure for 10-30 iterations. To execute these iterations quickly, we first perform a large-scale SVD factorization of the entire dataset to reduce each pixel’s fluorescence timecourse from tens or hundreds of thousands of frames to 1-2,000 SVD components, which retain most of the single-cell variance (Methods). This reduces the time required for each of the cell detection iterations to only a few seconds for standard recordings with hundreds to thousands of ROIs/plane.

When, as a control, we did not model the neuropil effectively, a large number of extracted ROIs ended up reflecting the neuropil, often at the expense of finding true cells (Supplementary Figure 2).

### Stage 3: ROI labelling and quality control

The outcome of stage 2 is a set of ROIs containing significant amounts of fluorescence variance (Figure 1c). Some of these active ROIs however might not be cells, but compartments such as dendrites or axons. Therefore, cell and non-cell ROIs must be distinguished for further analyses. These distinctions are first made by an automated classifier, and then curated by a human operator, using a GUI that displays information on each ROI and allows the classifier’s decisions to be overridden; finally, the classifier is retrained on the basis of these human decisions. This semi-automated process makes the quality control stage rapid, easy and intuitive.

The GUI shows large numbers of cells at the same time, allowing our visual system to do a parallel search for true negatives in the “rejected” view, and false positives in the “accepted” view (Figure 4). The GUI allows the user to rapidly switch between different views of the data, including mean image, variance image, pseudocolor pixel clustering image, red channel image, and several statistics computed by Suite2p (and used for automated curation, see below). The GUI can also display the timecourse of recorded fluorescence and estimated neuropil contamination for any selected ROI. Finally, the GUI allows the user to override automated ROI classifications either on an individual cell basis or globally (by adjusting the overall classification threshold).

**Figure 4.**
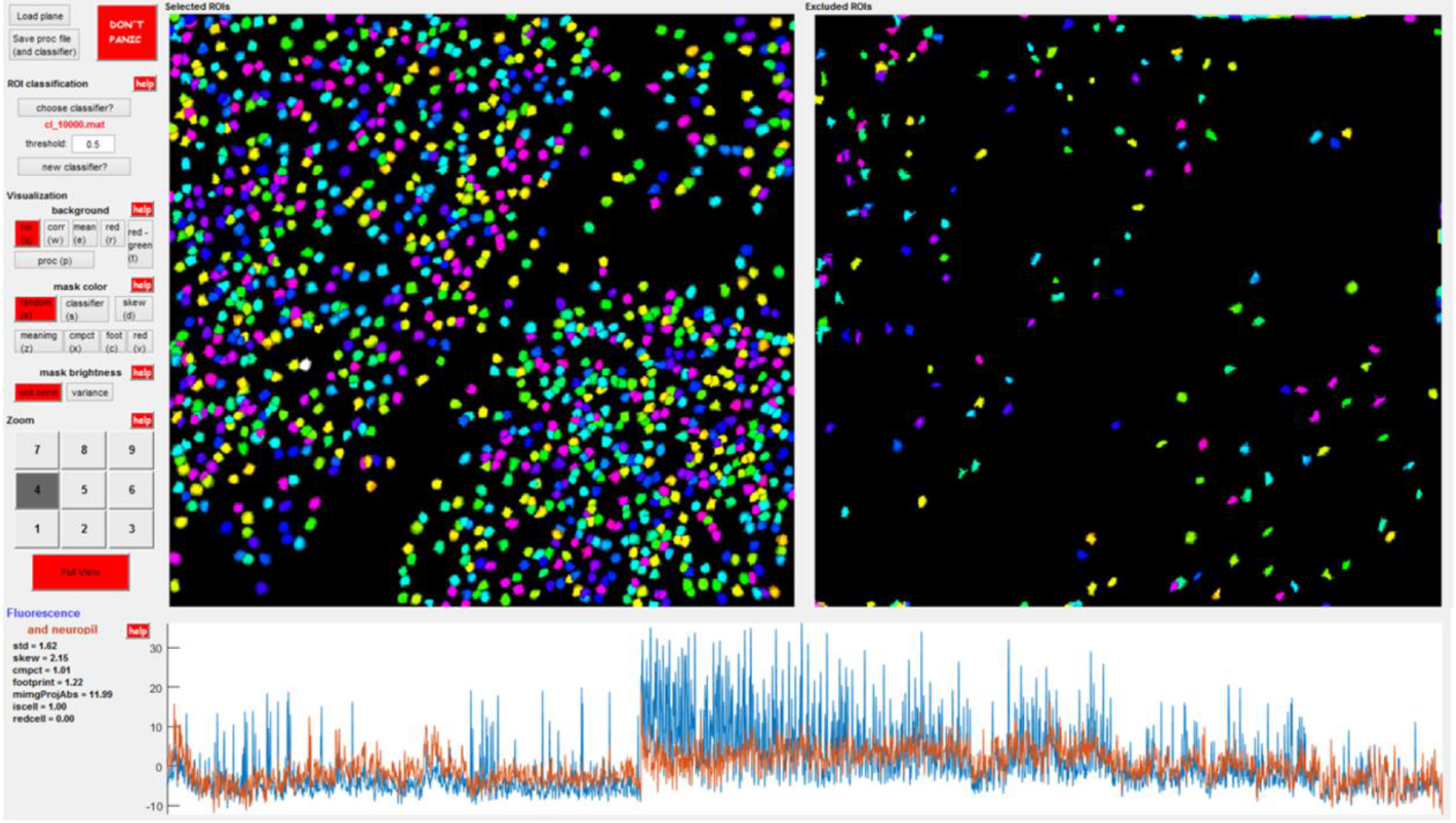
Screenshot of the graphical user interface used for quality control. The two pseudocolor panels show ROIs identified by Suite2P. The left panel shows the “accepted” ROIs, while the right panel shows the “rejected” ROIs. The GUI allows for curation of these automated results. The starting state of the GUI’s label assignments are provided by an automated classifier built based on multiple statistics available for each ROI, and a human curator can correct the classifier’s output. The classifier is updated after every manual session and improves based on the user’s choices. Several visualization methods are provided to enable users to quickly view the activity trace and other statistics of each ROI, to aid the manual decision. Detailed instructions for using the GUI are included online.

To improve manual curation, we developed a super-resolution method that increases the effective resolution of the mean image, allowing users to better infer the shapes of true cells. Our method takes advantage of the continuous brain motion generated by a headfixed animal. This motion generates XY displacement of the brain relative to the objective, which leads to frames being sampled at a continuous range of XY offsets. We can in turn effectively identify these offsets, using the subpixel registration method described in Stage 1, with a slight adjustment: instead of shifting frames by sub-integer amounts as we normally do, we bin frames by their XY offsets at subpixel resolution (25 bins, with 5 bins/pixel in X and 5 bins/pixel in Y). We then determine a mean image for each of the XY bins, and combine these mean images together into a super-resolution image. The final result adds substantial resolution, allowing the user to validate ROIs by their circular shape, which would not have been visible otherwise (Supplementary Figure 3). The super-resolution image is particularly useful in “zoomed out” recordings, where the resolution is too low to visually infer the shapes of all ROIs, leading to ambiguities during manual curation.

We also developed a GUI for matching cells found by Suite2p in separate recording sessions, for example across multiple days (Supplementary Figure 4). The GUI computes an affine transformation between the two recorded sessions based on user identification of matching ROIs, computes an optimal affine transformation based on these, and matches ROIs with substantial overlap after the transformation.

The classifier implemented in the GUI automatically labels ROIs as cells/non-cells, based on several statistics computed for each ROI. Some of these statistics are activity-dependent (skewness, variance, correlation to surrounding pixels), while others relate to the anatomical shape of the ROI (area, aspect ratio). The distribution of each statistic is estimated non-parametrically (see Methods) based on human-labelled data, and a naïve Bayes classifier based on these distributions is then applied to new data.

To account for different measurement statistics in different types of recordings, we allowed the classifier to be optimized separately for specific classes of dataset. The classifier substantially reduces the amount of human operator time required. For example, on large-scale data from primary visual cortex, the raw output of stage 2 cell detection contains 15% non-somas, as determined by a human curator (Supplementary Figure 5). After training the classifier, only 5% of the automatically-selected ROIs are not somas, at the cost of missing 1,5% of the somas identified by the human curator. The classifier also provides a quality metric, in the form of a posterior probability that each ROI represents a soma. This metric can be used to ensure consistency of subsequent analyses: for example, if a scientific result were to hold only for ROIs scoring low on this metric, that result might apply solely to non-somatic compartments.

### Stage 4: signal extraction and spike deconvolution

The final stage of the pipeline is to extract a single fluorescence signal for each ROI, representing that ROI’s calcium fluorescence timecourse. Crucially, this fluorescence signal needs to be corrected by removing, as well as possible, neuropil contamination signals. It may also be useful, in some circumstances, to estimate the spike train that underlies the measured calcium trace.

Our approach is to combine these steps – removal of neuropil contamination and estimation of underlying spike trains – by estimating parameters of a single generative model of the uncorrected fluorescence. Whether or not the users are interested in the estimated spike trains, this approach provides a robust method to remove neuropil contamination.

In the generative model, the uncorrected fluorescence **F_i_** is the sum of a somatic signal due to a spike train *s_i_*, a neuropil contribution **N_i_** scaled with a coefficient *c_i_*, and Gaussian noise:

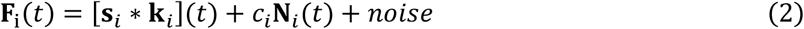

where ***s_i_*** * **k_*i*_** is the convolution of the spike train ***s_i_** > 0* with the calcium response kernel ***k_i_***.

To estimate this model, we start by extracting an uncorrected fluorescence signal **F_*i*_** for each ROI*i*. We do this by averaging pixels inside it, using the positive masks Λ*_ki_* from equation (1) as weights, and discarding any pixels belonging to multiple ROIs. We use this approach rather than the imputed source **f**_*i*_ found in the clustering step (equation 1), because the latter implicitly involves negative weighting of pixels, which carries a risk of obscuring problems in the datasets by constraining them to look like genuine spike trains (see Discussion).

Next, we compute the neuropil trace **N**_*i*_ for each ROI. Again, we avoid using the imputed neuropil traces ***n**_i_* found in the clustering step, as those traces implicitly use pixels inside ROIs and are not scaled by a cell-specific contamination coefficient. Rather, we average pixels in a “donut”-like area surrounding the ROI, following standard practice^3^. The neuropil trace **N**_i_ obtained in this manner is approximately linearly related to the expected neuropil contamination at the soma, but requires a further scaling coefficient *c*_i_. This scaling coefficient accounts for the fact that the ROI contains a cell that displaces some of the neuropil mass. Thus, the neuropil contamination coefficient must be less than 1. Two approaches have been suggested for setting this coefficient: a fixed value of 0.7 ^3,21^ and skewness maximization ^22^. The first ignores variations in coefficient across cells, while the latter might be a poor optimization criterion for classes of neurons that have inherently less skewed traces, such as interneurons ^23^, and can also return coefficients outside the physiological interval between 0 and 1 (our observations, not shown).

We propose a new approach, that estimates simultaneously a neuropil coefficient *c_i_* and a spike train *s_i_* for each cell *i*, by optimizing a cost function

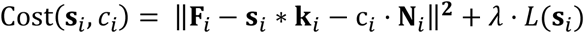

where *L*(s_*i*_) is an optional penalty on the spike train s_*i*_, whose entries are allowed to take continuous, non-negative values. To optimize this cost function, we iterate between optimizing *c_i_* while keeping s_*i*_ fixed, and optimizing s_*i*_ and k_*i*_ while keeping *c_i_* fixed. The first step corresponds to a simple regression problem, while the second step is a spike deconvolution routine appropriate for the penalty type *L*(s_*i*_) (*L_0_* norm, *L_1_* norm or no penalty).

The software allows several options for spike deconvolution. However, based on a quantitative comparison of multiple methods^24^, we recommend by default unconstrained non-negative deconvolution using exponential kernels ^25–27^ with a kernel decay timescale that matches the value reported in the literature for the particular calcium sensor used.

At the end of this stage, the user has obtained a neuropil-corrected calcium trace **F**_*i*_ − c__*i*__**N**_*i*_ for each ROI *i*. Moreover, if desired, the user has access to estimates of the spike times s_*i*_ in that ROI.

### Application to single and multi-plane data and benchmarks

To evaluate the performance of Suite2p, we obtained four standard-size datasets and two large-scale datasets in the primary visual cortex of awake mice. The standard-size datasets were obtained following viral injection of GCaMP6m, using conventional imaging methods: 5 plane imaging, ~500μm filed of view (FOV). The large-scale datasets, instead, were obtained in transgenic mice expressing GCaMP6s in all excitatory neurons, using a larger number of planes (11 planes) and a larger FOV (~900μm), yielding >1,000 cells per plane. A movie of the data obtained in one of these large-scale data sets is available (www.youtube.com/watch?v=xr-flH2Ow2Y).

For comparison, we also ran CNMF, a state-of-the art and commonly used integrated framework for denoising, deconvolution and demixing of calcium signals ^5^. In its default mode, however, running CNMF on even the standard datasets would require an unrealistic amount of RAM memory. To overcome this difficulty, we used a temporal subsampling approach ^28^. This allowed CNMF to run on datasets with up to 5,000 frames at 512×512 frame resolution, on a workstation with 64GB RAM. We perform the same temporal subsampling in Suite2p by default. We do not believe that temporal subsampling substantially impacted the cell detection performance of either algorithm, on these datasets. However, cell activity extraction in CNMF was severely compromised by subsampling, thus we did not evaluate that aspect of the pipeline here. Processing the large scale-recordings with CNMF was not possible on our workstations.

On temporally subsampled data, a single iteration of CNMF optimization took ~8 minutes, while a Suite2p iteration took ~7 seconds. This large difference in run time was due to Suite2p’s efficient source extraction algorithm, and due to the compact representation of the data provided by the SVD low-rank approximation. CNMF cannot use this low-rank approximation, because it would invalidate the positivity constraints that CNMF imposes on the calcium dynamics. In its default configuration, CNMF performs 2 iterations of optimization. Taking advantage of the decreased runtimes, Suite2p performs 10-30 iterations, until convergence.

To evaluate the accuracy of the algorithms, we compared the number of cells found by each. Because we had found that running Suite2p with a single global neuropil signal resulted in a large number of false ROIs to explain the neuropil’s spatial variation (Supplementary Figure 2), and because CNMF uses such a global neuropil model, we hypothesized that CNMF would also exhibit similar behavior. To test this, we ran both algorithms, setting their parameters so that the same total number of ROIs would be detected, and manually counted how many of those ROIs had soma-like morphology in the mean image, and calcium transients that were not visible in the surrounding neuropil (Figure 5a,b). Suite2p detected almost twice as many cells passing this criterion (Figure 5c). Furthermore, ROC analysis showed that the fraction of false-positive results was reliably lower with Suite2p than CNMF, when using a cutoff defined by the respective algorithms’ cell quality measures (Figure 5d). We also compared the average yields to an algorithm based on greedy segmentation of nearby pixels^1,8,12,13^, that we call “autoROI”. AutoROI required significantly more human curation, and found even fewer cells than CNMF, or about a third of the cells found by Suite2p (Supplementary Figure 6).

**Figure 5.**
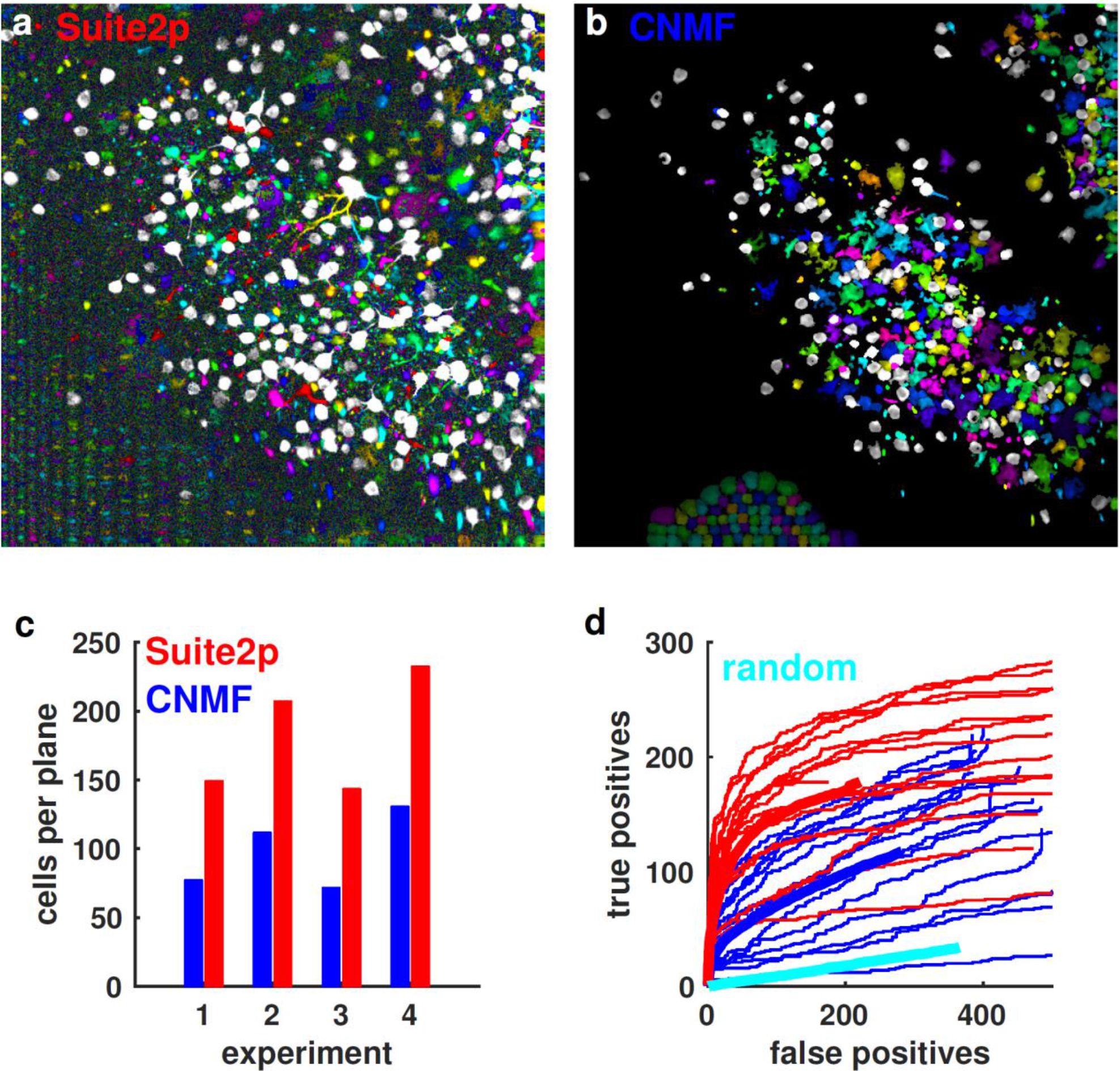
Detection performance compared to CNMF. (a) Results of Suite2p followed by manual curation. White ROIs were manually labelled as cells, while colored ROIs were rejected. (b) Same as (a) for CNMF. (c) Total cell yield comparison after manual curation on four datasets with four recorded planes each. (d) ROC curves showing the number of true versus false positives (defined by human curation), as increasing numbers of ROIs were accepted using the measure of cell quality produced by each algorithm. The averages for both methods, and for random selection of source centers, are shown in thick lines.

We finally turned to the large-scale datasets, and found that Suite2p obtained the activity of an unprecedented number of cells: over 10,000 (Figure 6). We were not able to run CNMF on these datasets, even with temporal subsampling, due to memory requirements larger than 64 GB. Suite2p, by contrast, was able to process the large datasets, thanks to its fast running speed and low memory demands. For example, the algorithm detected 13,451 cells in a 2 hour 11-plane imaging session with 900 μm field of view (Figure 6); for this dataset, the automated sorting took ~10 minutes/plane on a Core i7 workstation, 32 GB RAM and Nvidia GTX 980Ti GPU, and manual curation took an additional ~10 minutes/plane. The value of obtaining data at this scale can be further appreciated by observing robust and selective neural responses to stimuli on a trial-by-trial basis. We illustrate these responses for a ~7,000 neuron recording, by displaying the data in a manner similar to electrophysiology (“raster” plots, Supplementary Figure 7).

**Figure 6.**
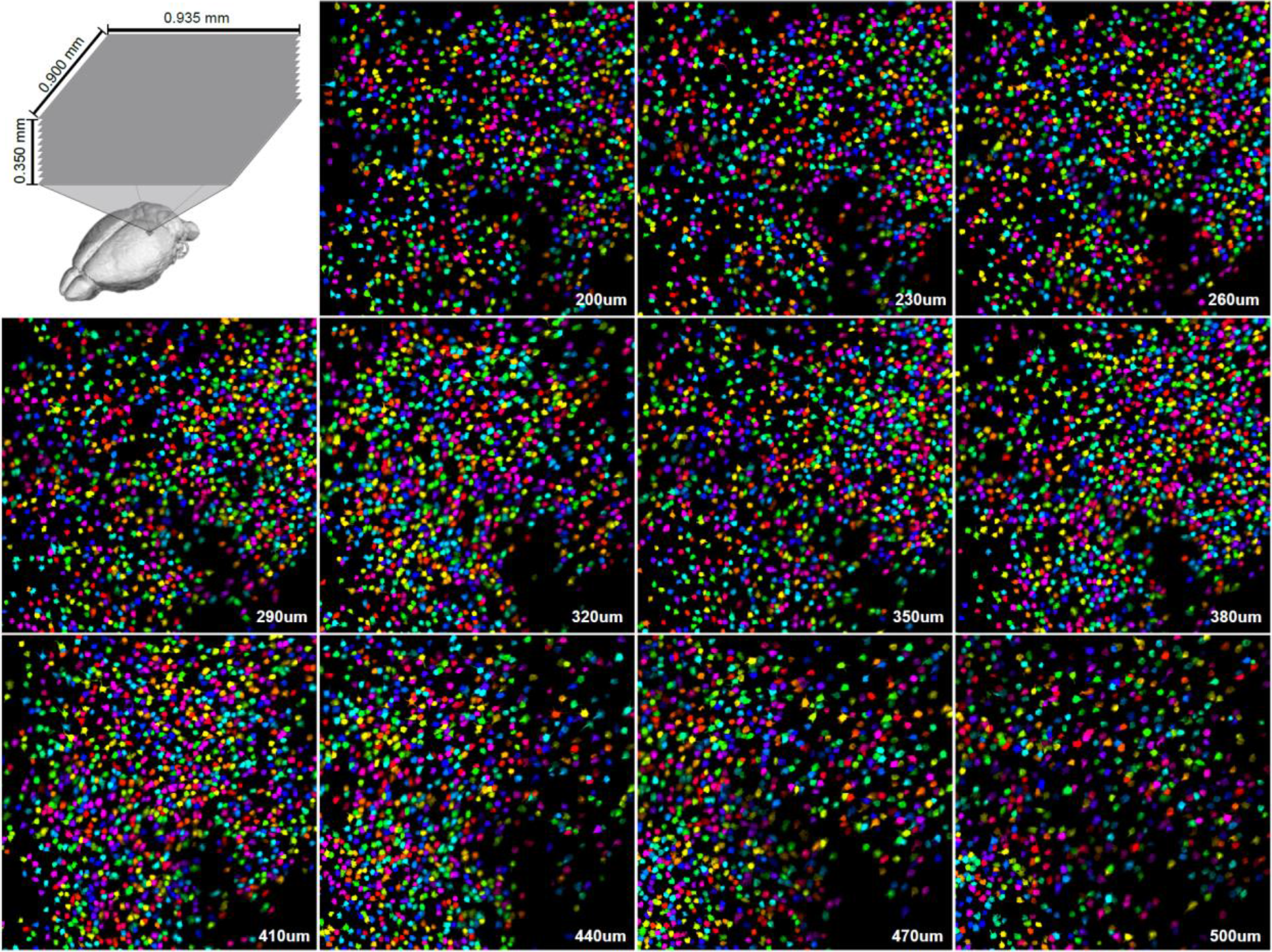
13,451 simultaneously recorded neurons. Imaging of an entire local cortical population, recorded from 11 planes in mouse visual cortex (GCaMP6s expressed in pyramidal cells of all layers using an Ai94; *Camk2a-tTA; Emx1-Cre* triple transgenic), using a standard Thorlabs 13-scope with resonant scanning at 2.5Hz frame rate. The pseudocolor masks of all 13,451 detected cells are shown. A short segment of the raw fluorescence in all planes can be seen at https://www.youtube.com/watch?v=xr-flH2Ow2Y.

### Application to dendritic spines and axonal boutons

Two-photon calcium imaging is not restricted to somata, but can also record the activity in neuronal subcellular compartments: dendrites, spines and axonal boutons, providing a powerful opportunity to study synaptic transmission and dendritic integration *in vivo* ^3^,^29^. However, because these compartments are smaller, and have different biophysical characteristics from neuronal somas, their accurate segmentation poses additional challenges.

To evaluate whether Suite2p could detect axonal and dendritic signals, we applied the algorithm to recordings from thalamic axonal boutons in the primary visual cortex (Figure 7a; injection at stereotaxic coordinates for LGN) and on recordings from sparsely labelled dendrites in L1 in the primary visual cortex (Figure 7c; injection of diluted AAV2/1-CAMK2a-Cre together with concentrated AAV2/1-hSyn-flex-GCaMP6 in V1 ^3^). Because Suite2p constrains ROIs to be local, we reasoned it should allocate separated boutons and spines to different ROIs, even though their time courses may be highly correlated. As expected, Suite2P allocated spines and boutons to different ROIs, even when belonging to the same parent dendrite or axon (Figure 7ac, insets) and extracted signals with differential timecourses (Figure 7bd). Suite2p, therefore, is suitable not only for analyzing two-photon data at the level of neuronal populations but also at the level of subcellular compartments such as spines and boutons.

**Figure 7.**
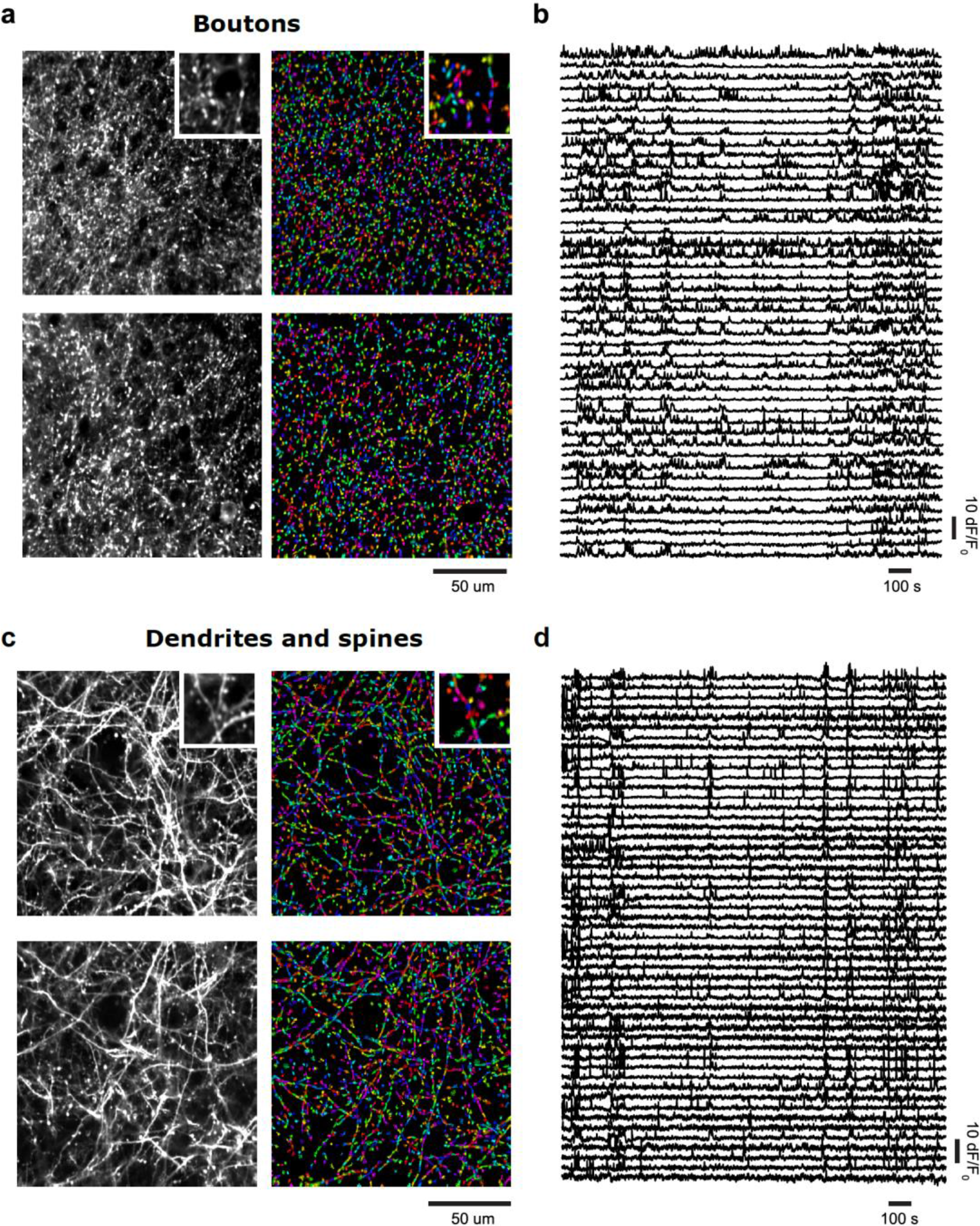
Detection of calcium transients in axonal boutons, and sparsely labelled dendrites. (a) Left panels: average projections of registration output from recordings of two axonal FOVs acquired simultaneously at high magnification, at 20pm vertical spacing. Neurons in lateral geniculate nucleus were virally labelled with GCaMP6s, and their axons imaged in cortical layer 4. Right panels: ROI segmentation of the axonal fields identifies active boutons. Insets: magnified by factor of 2. (b) GCaMP6s trace extracted by Suite2p from some example boutons. (c) Left panels: average projections of registration output from the recordings of two fields of view in cortical layer 1 acquired simultaneously at high magnification, at 20pm vertical spacing. Dendrites were labelled with GCaMP6s and tdTomato (see Methods). Right panels: Suite2p ROI segmentation of the dendrites fields identifies active dendrites and spines. Insets: magnified by factor of 2. (d) GCaMP6s trace extracted by Suite2p from some example dendrites and spines.

## Discussion

Suite2p is a fast and accurate set of tools for processing of two photon calcium imaging data, allowing detection of over 10,000 cells in multi-plane recordings. The pipeline is modular, allowing the main processing steps to be used independently: registration, cell detection, spike deconvolution and manual GUI.

One of the central goals in the development of Suite2p has been to allow rapid curation and quality control of all processing steps, so that any problems with data quality can be quickly recognized. As well its built-in GUI, Suite2p achieves this goal by reducing the number of free parameters, and avoiding algorithmic constraints that would force even poor quality data to look like genuine neural activity. For example, the activity traces we compute are simple weighted averages of raw pixel recordings; neuropil contamination is rejected by subtraction of pixels surrounding the ROI; and our default spike deconvolution method does not assume any sparsity penalties on the extracted spikes (L0 or L1) or attempt to infer optimal temporal kernels for each cell. Indeed, introducing these complexities into the algorithm actually reduces spike estimation performance on available ground truth data ^24^.

One of the most difficult challenges facing densely labelled calcium imaging is neuropil (or background) contamination. Neuropil activity is not random noise, but can contain responses to sensory stimuli ^20^: failure to decontaminate the activity trace of a cell can therefore lead to invalid scientific conclusions, by biasing the tuning of that cell towards the tuning of the neuropil ^30^. We found that modeling the spatial distribution of the neuropil provided substantial performance improvement in the ROI detection step by enabling dimmer cells to be distinguished from the surrounding background (Figure 3, Supplementary Figure 2).

Previous approaches to this problem have employed matrix decomposition procedures, while simultaneously enforcing constraints on the extracted traces, such as sparsity, non-negativity, or auto-regressive dynamics ^5,7,9^. Typically, these approaches allow pixels to contribute positively or negatively to an inferred ROI’s fluorescence, and can even extract ROI fluorescence as a nonlinear combination of pixels. While this approach in principle allows for automatic subtraction of contamination by neuropil signals and other cells, it carries substantial risks. Indeed, by constraining the extracted signals to look like genuine calcium traces, such flexible methods may produce apparently high-quality traces despite problems in the original data such as neuropil contamination, virus overexpression, and contamination by hemodynamic or light sources. Simple extraction as pixel averages would reveal these problems, but additional constrained processing would incorrectly “fix” them, forcing the extracted traces to have the properties of genuine calcium traces even when underlying data cannot be salvaged.

Our results show that constrained extraction is not necessary, and that it is safer to remove neuropil contamination using specifically tailored methods. Indeed, even in densely-labeled neocortical volumes, overlap between cells is minimal, as the distance between cellular somata is generally larger than the point-spread function even in the Z-direction.

As well as somata, Suite2p is able to detect the activity of individual axonal boutons, dendritic compartments, and spines. Importantly, even though multiple terminals of a single axon or dendrite will have highly correlated activity, Suite2p’s spatial constraints ensure they are detected individually rather than as a single multi-location ROI. This will enable analyses of differential activity such as synaptic inputs to individual spines, integration in dendrites and the probability of release from different boutons (Figure 8AB). If required, identification of ‘sibling’ dendrites, spines and boutons can be achieved post-hoc by means of signal correlation or morphological tracing.

One way to compare the performance of different analysis methods is through open competitions, where different groups can test their algorithms on publicly available data, and the organizers hold the results of ground-truth measurements aside to evaluate and rank the different analyses. An example of such a competition at the time of writing is the Neurofinder challenge, where Suite2p scored a high value of 0.571, close to the 0.597 score of the top ranking algorithm HNCcorr ^31^. However, these results should be interpreted with caution, because in this challenge the “ground truth” labels are provided by human curators; if these curators miss some cells, an algorithm will be incorrectly penalized for finding them. Indeed, visual inspection of the ROIs detected by Suite2p but not by Neurofinder’s human curators suggests that it found rarely-firing cells missed by human operators in these relatively short (5-10 minute) datasets. Therefore, Suite2p’s performance may in fact be substantially greater than Neurofinder’s estimates.

Suite2p enables routine recordings of more than 10,000 cells. We have successfully used the tools presented here to scale up the yields of two-photon recordings and automate most of the processing of ~10,000 neurons recorded simultaneously at 3Hz sampling rates (Figure 6). This has now become a routine experiment in our laboratory, but was not practical using existing processing pipelines ^5,7,9^ that took days or weeks of computation to transform these data into single-neuron activity traces or spikes. Although a case has been made for parallelizing such pipelines over large compute clusters ^4,32^, such computing resources are not available to many neuroscience laboratories except via expensive commercial rental. Furthermore, we have also shown that Suite2p outperforms the state-of-the-art algorithm (CNMF) in terms of the neural yields after manual curation, by a factor of approximately 2.

The large yields and high SNR of our large-scale recordings were enabled not just by the efficiency of Suite2p, but also by other considerations. Specifically, we used the GCaMP6 slow sensor for its high SNR, rather than the fast version, because theoretical considerations show it to have a better relationship to ground truth spiking ^33^, because its high SNR allows a much larger fraction of all neurons to be detected ^3^, and because its slow temporal dynamics allow the use of 3Hz scan rates without the danger of missing action potentials between frames. We also used high laser powers (≈ 100mW), though still well below the limit of 200mW for avoiding heat damage, calculated by Ref. ^34^. Using these recording protocols in combination with efficient automated processing methods, will allow the characterization of complete neural populations on a single-trial basis.

## Materials and methods

### Imaging in visual cortex

All experimental procedures were conducted according to the UK Animals Scientific Procedures Act (1986). Experiments were performed at University College London under personal and project licenses released by the Home Office following appropriate ethics review.

The experimental methods were similar to those described elsewhere (Dipoppa et al, 2016). Briefly, surgeries were performed in adult mice (P35–P125) of multiple lines (wild type C56bl6/j, Ai94;CamK2a-tta;Emx1-Cre triple transgenics, and Gad-Cre transgenics) in a stereotaxic frame and under isoflurane anesthesia (5% for induction, 0.5-1% during the surgery). During the surgery we implanted a head-plate for later head-fixation, made a craniotomy with a cranial window implant for optical access, and, on relevant experiments, performed virus injections with a beveled micropipette using a Nanoject II injector (Drummond Scientific Company, Broomall, PA 1) attached to a stereotaxic micromanipulator. Depending on the experiment, we used the following viruses: AAV2/1-CamKII-Cre, AAV2/1-hSyn-flex-GCaMP6s, AAV2/1-CAG-flex-tdTomato, AAV2/1-hSyn-GCaMP6m/s. Viruses were acquired from University of Pennsylvania Viral Vector Core. Injections of 50-200 nl virus (1-3 x10^12^ GC/ml) were targeted to monocular V1, 2.1-3.3 mm laterally and 3.5-4.0mm posteriorly from Bregma and at a depth of L2/3 (200-400 μm).

### Hardware

We tested Suite2p on PC systems with Intel i7 processors, 16,32 or 64 GB of RAM, 256GB SATA SSD or 512GB PCIe SSD, and one of GTX 970, 980Ti, 1060, 1070 or 1080 GPUs. Systems with these specifications can be bought for $1000-$2,000.

### CNMF implementation

To test the CNMF pipeline ^5^ we used public code from https://github.com/epnev/ca source extraction. We also used OASIS ^25^, a fast spike deconvolution with L1 penalties https://github.com/zhoupc/OASIS matlab. We provide wrapper functions in Suite2p for using OASIS without an L1 penalty, together with the neuropil estimation step described in the paper, and with the baselining and pre-processing routines of Suite2p.

### Accelerating cell detection with large-scale SVD decomposition

The execution speed of the algorithm is greatly boosted by a simple computational trick. After spatial registration, we reduce the dimensionality of the data using an approximate singular value decomposition (SVD), and perform all optimization iterations in a space of much lower dimension than the original movies. This step greatly reduces the space required to store the data, denoises the data, and allows our modelling framework to operate fast enough for large-scale recordings (10-100x depending on original number of frames in the dataset). To perform this optimizaiton, we replace the temporal activity of each pixel *k* with a vector of SVD coefficients **U**_*k*_. We obtain these coefficients by approximating the recorded fluorescence ***F*** (represented as a matrix with one row for each pixel and one column for each timepoint) of all pixels *k* at times *t* with its best approximation ***F** ~ **U V***^T^, where ***U*** has as many rows as pixels, ***V*** has as many rows as timepoints, and both have a limited number of columns. The singular values have been absorbed into ***U***, which is an orthogonal but not orthonormal matrix, while ***V*** is an orthonormal matrix.

We use an approximate SVD because the full SVD would require unrealistic running time on the large-scale data we acquire. To do this, we temporally bin the signal at each pixel until the number of timepoints is reduced to under 10,000. We then compute the top eigenvectors **V** of the time-by-time covariance matrix (1,000 by default), and use these to compute ***U = F V***. Once cell masks and neuropil weightings have been estimated, we compute the full fluorescence timecourses of each cell using the raw, non-binned data. This downsampling approach has similarities to another recent approach ^28^, but the authors of that study also propose to spatially downsample the data. While such an approach would indeed save computational costs, it loses resolution for resolving and distinguishing ROIs, and thus requires a high magnification recording. In contrast, the SVD decomposition used here retains fine spatial features and allows even greater computational acceleration.

### ROI detection

Cell detection consists of optimizing the model described in the main text (Equation 1)

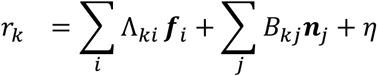

To fit this model, we alternate between three steps:

- Source detection. This step finds new sources, not yet accounted for. To find the new sources we analyze the residuals after subtracting off the current model, and identify locations where the residuals are highly correlated in a small spatial neighborhood. Specifically, we compute the “multi-dimensional correlation” of each pixel trace *r_k_* with pixel traces in its local neighborhood (see next section for more details). The peaks of this correlation map are putative new sources, and we select those that dominate their local neighborhood over a radius of at least twice the average cell radius. Each source introduced in this fashion is initialized at the corresponding location, and assigned a spatial mask Λ_*kn*_ given by the correlation of the central pixel with the neighboring pixels. Peaks that are not added on a given iteration because of a larger peak nearby will still be added in a future iteration.
- Source extraction. This step re-estimates the timecourses ***f**_i_* of all sources, as well as the timecourses ***n**_j_* of the neuropil, given the spatial masks Λ_*ki*_ of all sources and of the neuropil basis functions. This is achieved by minimizing the square of the error terms η. This operation amounts to a pseudo-inverse matrix computation.
- Source re-assignment. This step re-estimates the spatial distribution of the sources Λ*_kn_*. The spatial mask of each source is refined in turn, fixing the timecourses of all sources and the spatial masks of all other sources, and optimizing just the spatial mask of the considered source. Again the squared error is minimized, on each pixel individually. Pixel weights below a fifth of the maximum weight are replaced with zeros. At each step, we loop through all sources once in the order they have been added, and perform this operation independently for each. This step resembles the parameter optimization step of the K-SVD algorithm ^35^.

### Computing the correlation map for finding cells

To find the locations of new sources we compute the “multi-dimensional” correlation of each pixel with the surrounding pixels. We define the multi-dimensional correlation between a set of *N* traces ***f**_i_* (all with mean 0) as

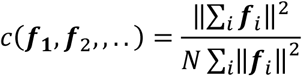

The multi-dimensional correlation is always bounded between 0 and 1, and is 1 only when ***f**_i_* = ***f**_j_* for all *i,j*. In general, the multi-dimensional correlation is high if and only if all the vectors ***f**_i_* point in approximately the same direction. We can also compute a weighted multi-dimensional correlation

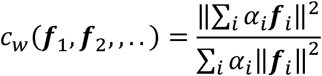

The weights can be, for example, a Gaussian-shaped kernel around each pixel, while ***f**_i_* are the respective traces of neighboring pixels. *c_w_* defined in this manner can be computed efficiently at each pixel, by smoothing with a Gaussian kernel each dimension of ***f**_i_*. The resulting “correlation maps” then contain the weighted version of this multi-dimensional correlation at each pixel location. The relatively large values in this map are putative locations for new sources. In practice, we also found it useful to top-hat filter the correlation maps locally, by subtracting off the morphological opening (of twice the expected cell diameter) from the raw map.

### Automatically labelling ROIs as somata with customizable classifiers

To automatically classify which ROIs are cells, we defined a set of *k* statistics *r_k_*(*n*) measured on each ROI *n*, and trained a classifier to reproduce the results of human curation on a training dataset. We found that linear classifiers were not suitable: for example, cells typically have an intermediate size, with both very small and very large ROIs being predominantly not cells. We therefore employed a simple nonlinear “naïve Bayes” classifier. For each statistic *x*, we estimate the positive and negative distributions 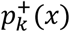 and 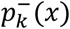 (i.e. the distributions classified by the operator as somata and non-somata) by adaptively smoothing their empirical histograms (smoothing width in each bin inversely proportional to number of samples in that bin). To automatically classify a new ROI *N*, we compare the scores 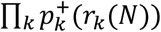 and 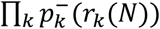, and assign the ROI to the category that gives it a larger score.

### Neurofinder benchmark

The neurofinder benchmark consists of 9 test datasets, using data from several laboratories, which used different annotation methods. We excluded datasets 1,2 and 8 from our comparison to HNCcorr ^31^, because on these datasets, anatomical algorithms perform significantly better, and the manual annotation did not use activity at all.

## Acknowledgements

We thank Charu Reddy for assistance with surgeries and Michael Krumin for assistance with the two-photon microscopes. This work was supported by the Wellcome Trust (95668, 95669, 108726), and the Simons Foundation (325512). CS was funded by a four-year Gatsby Foundation PhD studentship. MD and SS were supported by Marie Skłodowska-Curie Fellowships. LFR was funded by a four-year Wellcome Trust PhD studentship in Neuroscience. MC holds the GlaxoSmithKline / Fight for Sight Chair in Visual Neuroscience.

---

**Supplementary Figure 1.**
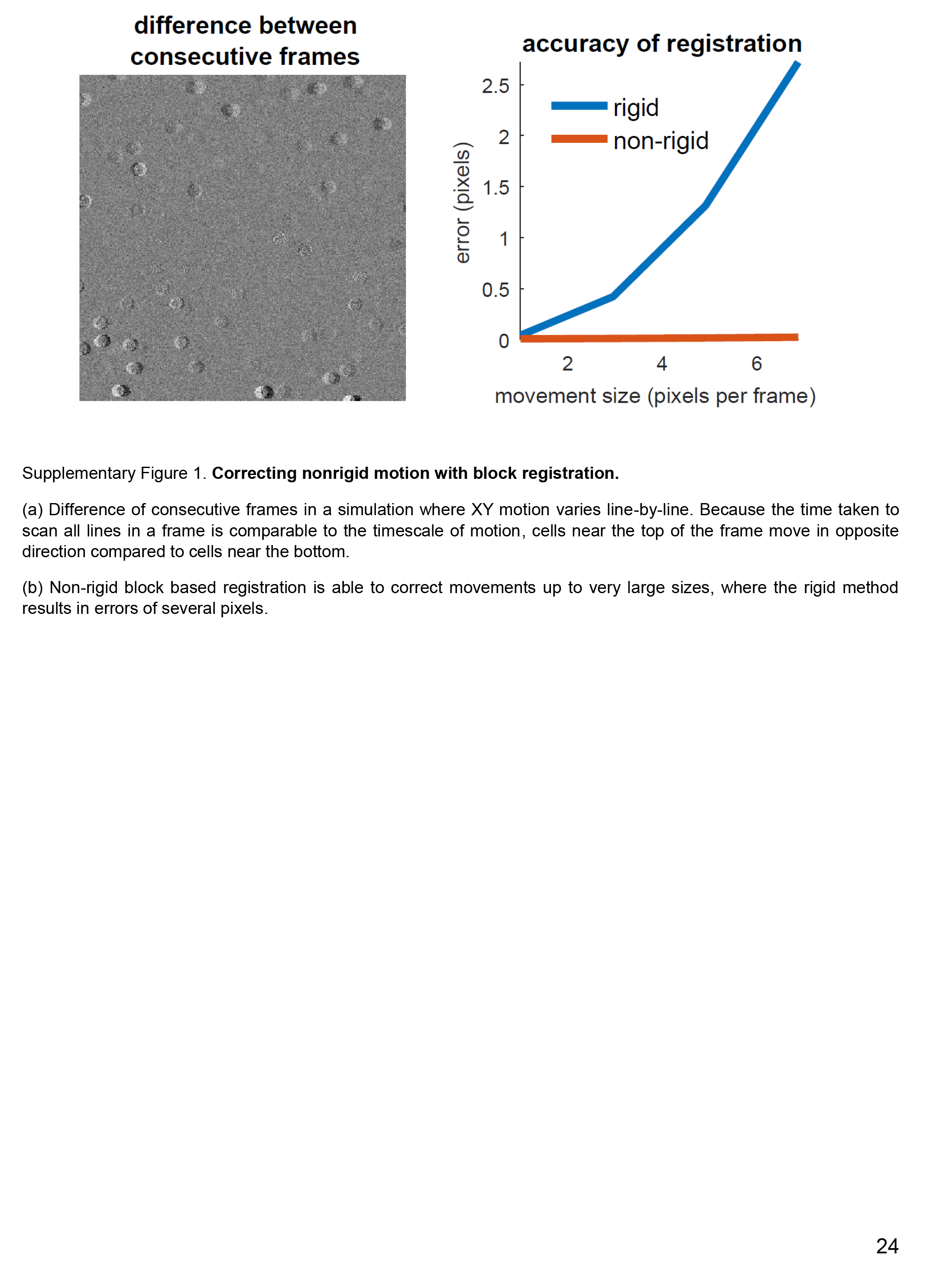
Correcting nonrigid motion with block registration. (a) Difference of consecutive frames in a simulation where XY motion varies line-by-line. Because the time taken to scan all lines in a frame is comparable to the timescale of motion, cells near the top of the frame move in opposite direction compared to cells near the bottom. (b) Non-rigid block based registration is able to correct movements up to very large sizes, where the rigid method results in errors of several pixels.

**Supplementary Figure 2.**
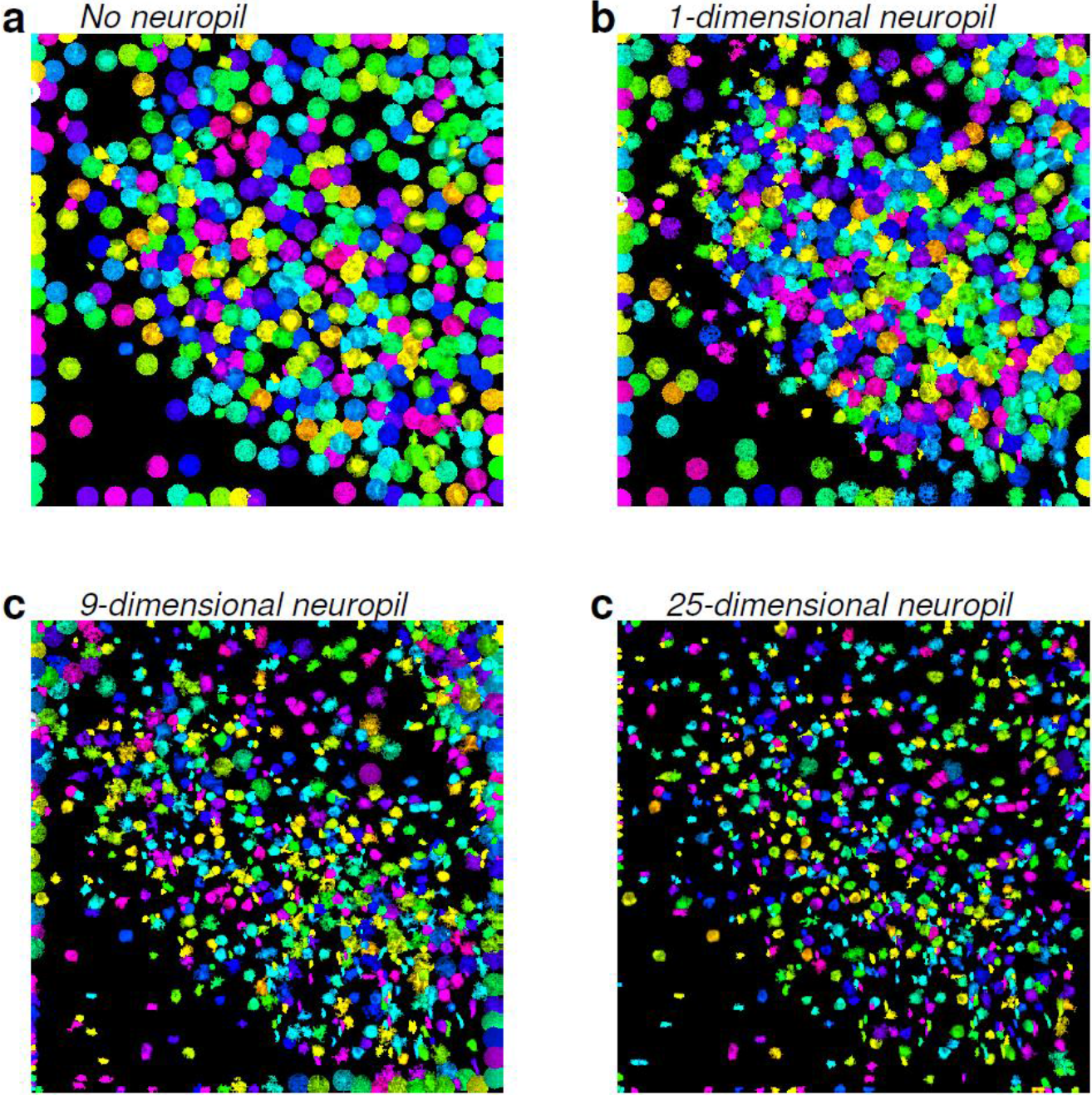
Modelling the spatial structure of neuropil contamination results in better-defined cellular ROIs. All ROIs found by Suite2p are shown, before the quality control step. Each ROI is assigned a random color. (a) Without a neuropil model, most ROIs are assigned to neuropil. (b) With a single, global neuropil component (i.e. all pixels receive same contribution), most ROIs are still assigned to neuropil, but some of the ROIs inferred are cells. (c) With 9 neuropil basis functions, most of the ROIs resemble cellular compartments, but ROIs close to the edge may not. (d) With 25 neuropil basis functions, almost all ROIs inferred appear to identify cellular compartments (somas or dendrites). This corresponds to the default setting in Suite2p, which uses basis functions of width six times the user-specified average cell diameter, 12 pixels for this recording.

**Supplementary Figure 3.**
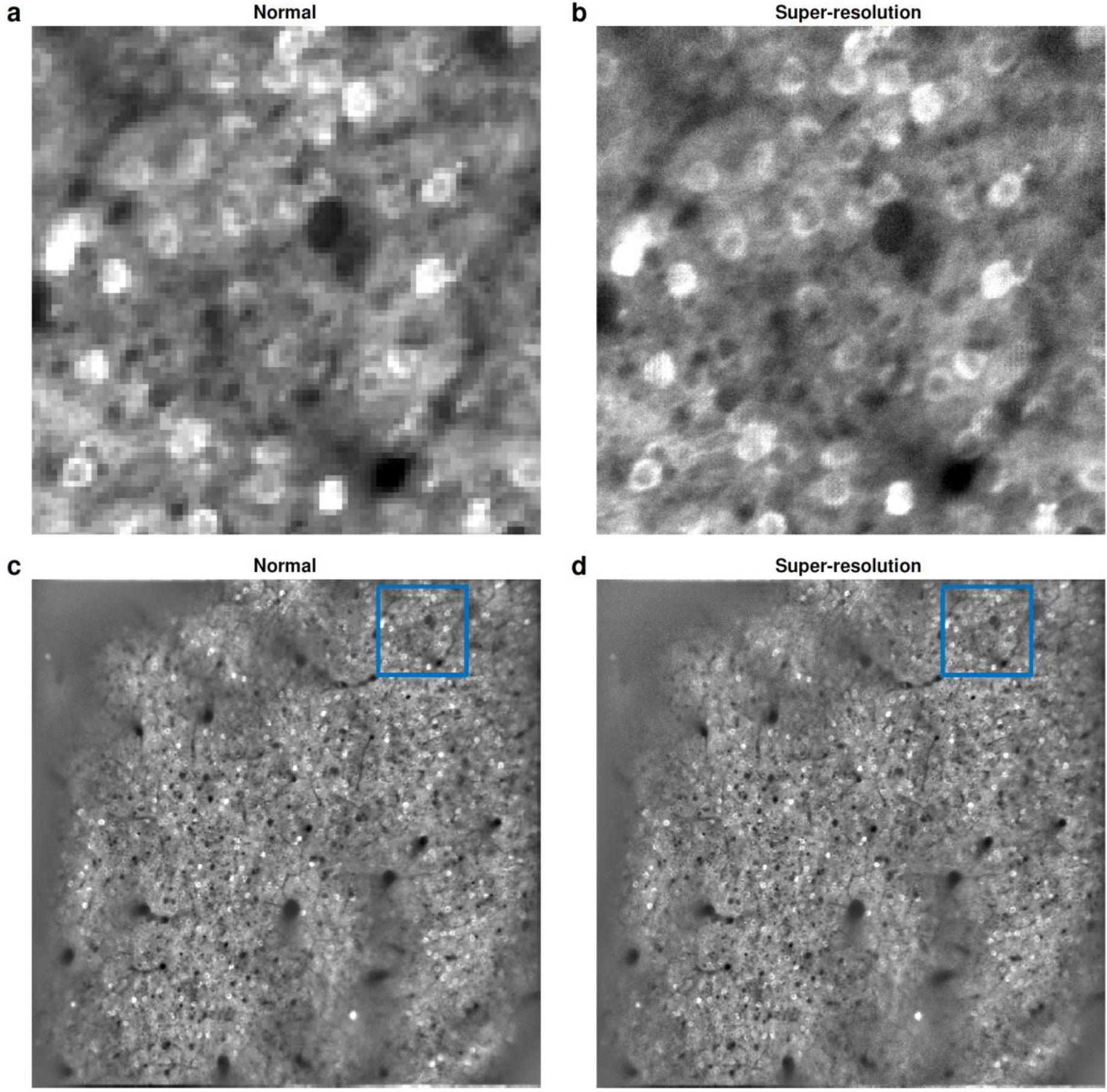
Super-resolution mean image. (a) Portion of the average image from a recording in V1 (Ai94; *Camk2a-tTA; Emx1-Cre* triple transgenic that expresses GCaMP6s in pyramidal cells of all layers). (b) Super-resolution version of the same field of view. (cd) Full field of view for original, and super-resolution mean image. The blue rectangles show the field of view from **ab**.

**Supplementary Figure 4.**
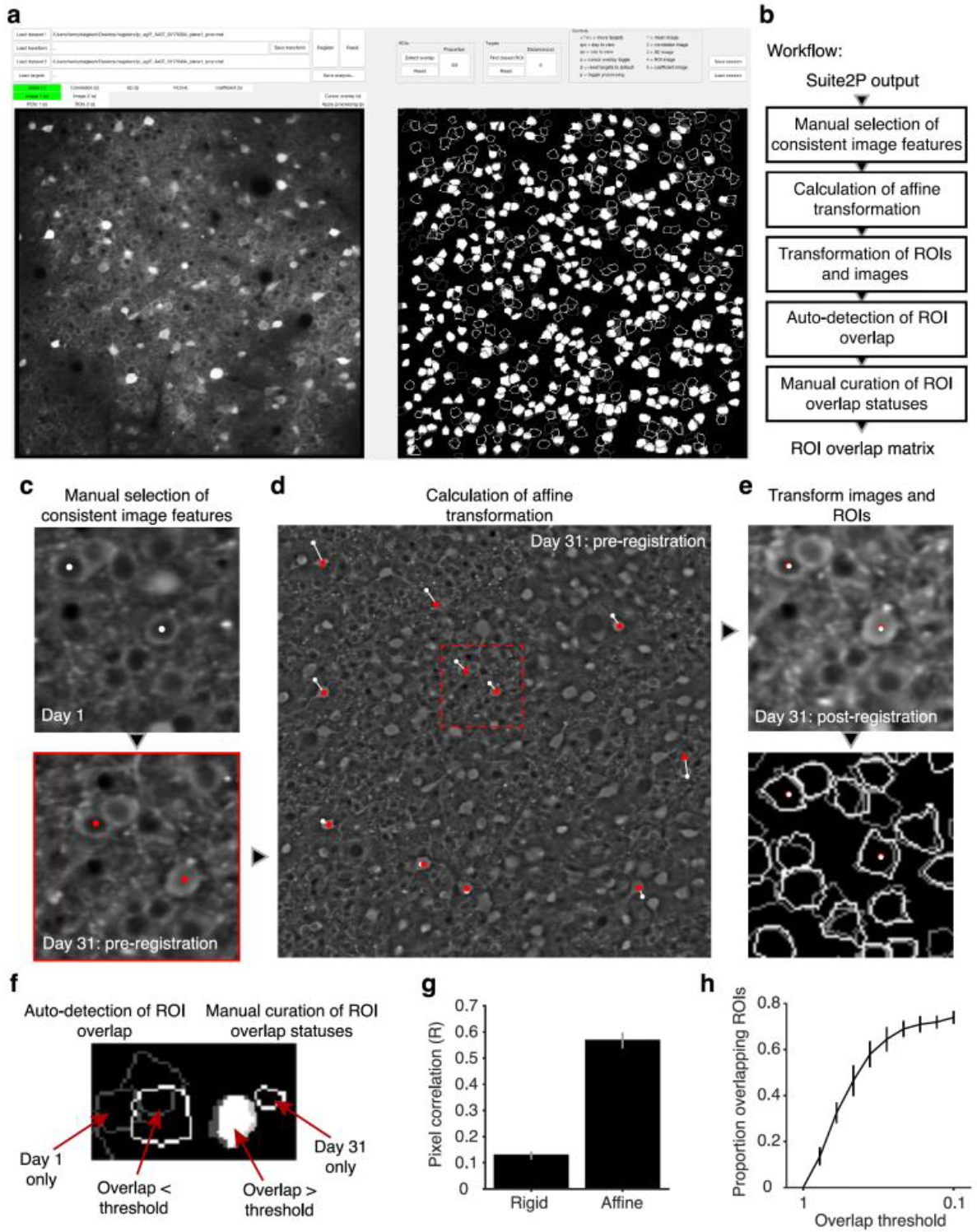
Semi-automatic ROI registration across days. (a) Graphical user interface showing FOV images (left; the user switches between the two sessions with a keypress) and ROIs (right, white and gray outlines for separate sessions; filled ROIs indicate overlaps). Datasets have been registered and overlapping ROIs have been automatically matched and manually validated (filled ROIs in right panel). (b) GUI workflow schematic. A user selects features that are then used to align the two sets of (c) Zoom-ins from an FOV imaged on day 1 (top) and day 31 pre-registration (bottom). A subset of consistent features are indicated with colored dots. Note shifts between the two images. The displayed images are spatially filtered (as in the normal Suite2p GUI) to reveal fine details. (d) Consistent features (white = day 1, red = day 31) are plotted on the mean image from day 31 with displacement vectors as white lines between them. These paired control points are used to calculate an affine transformation to register the two datasets. (e) Following registration, cellular features (top) and Suite2P ROIs (bottom) from day 31 overlap with those of day 1. Top: coordinates of day 1 features are plotted in white over the features from day 31 in red. Bottom: white contours = outlines of day 31 ROIs, grey contours = outlines of day 1 ROIs. (f) GUI representation of different overlap possibilities, auto-detected with a threshold of 60% of pixels. White: day 31, gray: day 1, filled: overlap > threshold, empty: overlap < threshold. ROI states can be manually toggled. (g) Affine transformations yield a significantly higher pixel-wise correlation of FOV images across time-points than rigid transformations (p = 0.006). (h) Proportion of day 31 ROIs that overlap with a day 1 ROI for different overlap thresholds. All n = 2 FOVs across 2 mice, all errorbars = SEM.

**Supplementary Figure 5.**
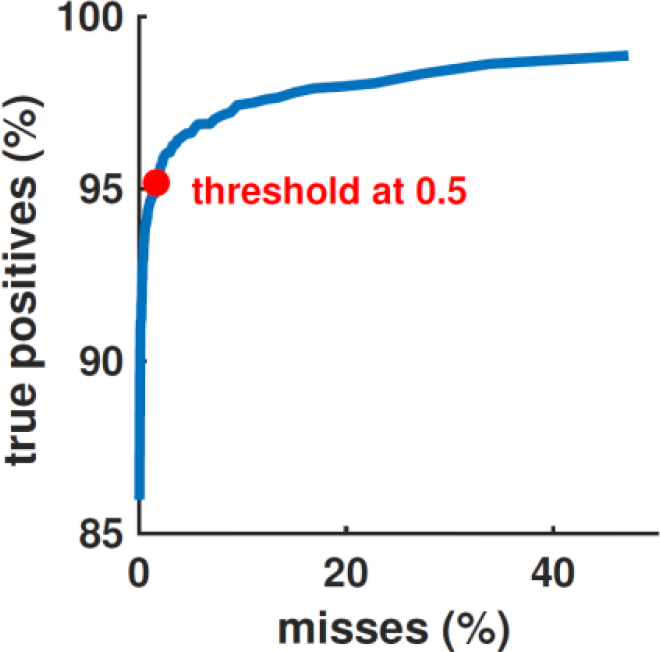
Improving quality control with customized classifiers. To further refine the output of Suite2p, we provide the option to train a classifier, based on the user’s own curated datasets. For the large-scale −10,000 cell recordings, such a classifier improves the rate of true positives (defined by human manual curation) from 85% to 95%, with only 1.5% misses. The classifier’s output is a continuous probability between 0 and 1, which can be used as a quality metric. By this metric, higher acceptance thresholds can further increase the rate of true positives to 99%, but at the cost of almost 50% misses.

**Supplementary Figure 6.**
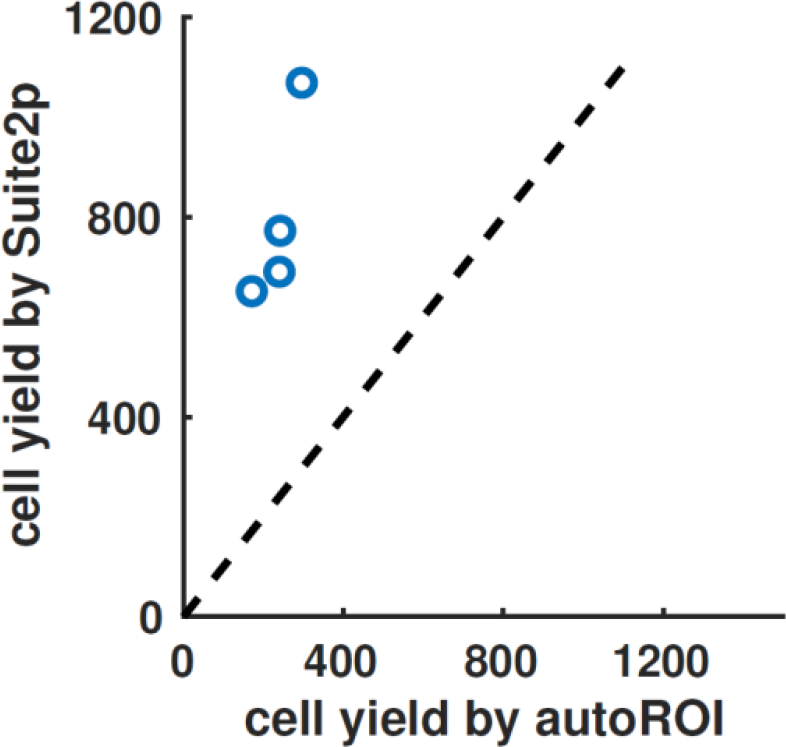
Cell yields using greedy pixel segmentation. We evaluated the performance of a method previously used in our lab (autoROI), which is based on greedy segmentations of correlated pixels. A human operator judged whether ROIs found by the method corresponded to genuine cells, similarly of the Suite2p output. The four datasets we analyzed were recorded in visual cortex at a spatial resolution of 512×512 over a 500μm × 500μm field of view, at 5Hz scan rate, using the GCaMP6m calcium sensor (see Methods). Suite2p found approximately three times more neurons, in each of the four recordings (792 vs 241 cells on average).

**Supplementary Figure 7.**
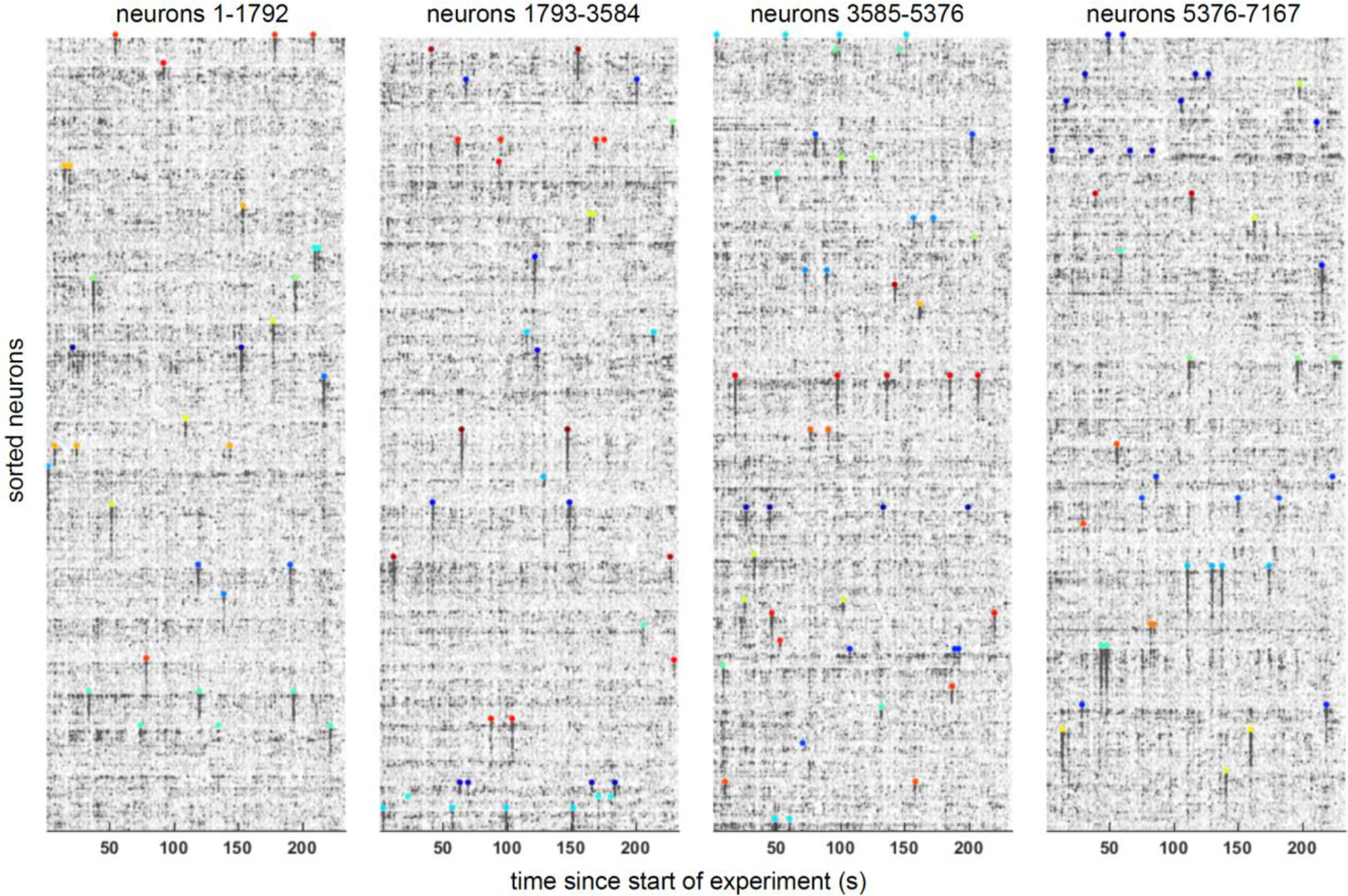
Spike deconvolution and responses to visual stimuli in 7,167 simultaneously-recorded neurons. We recorded nine planes from layer 2/3 of mouse visual cortex in a transgenic GCaMP6s-expressing line (Camk2a-tTA; Ai94(TITL-GCaMP6s); Rasgrf-Cre). We used a spike deconvolution method to obtain the most likely times of spikes and their burst magnitudes, and sorted all neurons by their preferred stimulus out of 112 natural images. The raster shows the deconvolved activity of all neurons for a duration of approx 4 minutes,. Colored dots represent stimulus times, located above the population of neurons which responded best to that stimulus.

## References

1. Sofroniew, N. J., Flickinger, D., King, J. & Svoboda, K. A large field of view two-photon mesoscope with subcellular resolution for in vivo imaging. bioRxiv 055947 (2016).

2. Stirman, J., Smith, I., Kudenov, M. & Smith, S. Wide Field-Of-View, Multi-Region Two-Photon Imaging Of Neuronal Activity In Vivo. in Optics and the Brain BTu2D-2 (Optical Society of America, 2016).

3. Chen, T.-W. et al. Ultrasensitive fluorescent proteins for imaging neuronal activity. Nature 499, 295–300 (2013).

4. Freeman, J. et al. Mapping brain activity at scale with cluster computing. Nat. Methods 11, 941–950 (2014).

5. Pnevmatikakis, E. A. et al. Simultaneous denoising, deconvolution, and demixing of calcium imaging data. Neuron (2016).

6. Romano, S. A. et al. A computational toolbox and step-by-step tutorial for the analysis of neuronal population dynamics in calcium imaging data. bioRxiv 103879 (2017).

7. Mukamel, E. A., Nimmerjahn, A. & Schnitzer, M. J. Automated Analysis of Cellular Signals from Large-Scale Calcium Imaging Data. Neuron 63, 747–760 (2009).

8. Kaifosh, P., Zaremba, J. D., Danielson, N. B. & Losonczy, A. SIMA: Python software for analysis of dynamic fluorescence imaging data. Front. Neuroinformatics 8, (2014).

9. Andilla, F. D. & Hamprecht, F. A. Sparse space-time deconvolution for calcium image analysis. in Advances in Neural Information Processing Systems 64–72 (2014).

10. Apthorpe, N. et al. Automatic Neuron Detection in Calcium Imaging Data Using Convolutional Networks. in Advances In Neural Information Processing Systems 3270–3278 (2016).

11. Pachitariu, M. et al. Extracting regions of interest from biological images with convolutional sparse block coding. in Advances in Neural Information Processing Systems 1745–1753 (2013).

12. Smith, S. L. & Hausser, M. Parallel processing of visual space by neighboring neurons in mouse visual cortex. Nat. Neurosci. 13, 1144–1149 (2010).

13. Vladimirov, N. et al. Light-sheet functional imaging in fictively behaving zebrafish. Nat. Methods (2014).

14. Poort, J. et al. Learning enhances sensory and multiple non-sensory representations in primary visual cortex. Neuron 86, 1478–1490 (2015).

15. Alba, A., Vigueras-Gomez, J. F., Arce-Santana, E. R. & Aguilar-Ponce, R. M. Phase correlation with sub-pixel accuracy: a comparative study in 1D and 2D. Comput. Vis. Image Underst. 137, 76–87 (2015).

16. Foroosh, H., Zerubia, J. B. & Berthod, M. Extension of phase correlation to subpixel registration. Image Process. IEEE Trans. On 11, 188–200 (2002).

17. Greenberg, D. S. & Kerr, J. N. Automated correction of fast motion artifacts for two-photon imaging of awake animals. J. Neurosci. Methods 176, 1–15 (2009).

18. Pnevmatikakis, E. A. & Giovannucci, A. NoRMCorre: An online algorithm for piecewise rigid motion correction of calcium imaging data. bioRxiv 108514 (2017).

19. Pachitariu, M. et al. Suite2p: beyond 10,000 neurons with standard two-photon microscopy. bioRxiv 061507 (2016).

20. Lee, S., Meyer, J. & Smirnakis, S. Visually driven neuropil activity and information encoding in mouse area V1. bioRxiv 113019 (2017).

21. Peron, S. P., Freeman, J., Iyer, V., Guo, C. & Svoboda, K. A cellular resolution map of barrel cortex activity during tactile behavior. Neuron 86, 783–799 (2015).

22. Bonin, V., Histed, M. H., Yurgenson, S. & Reid, R. C. Local diversity and fine-scale organization of receptive fields in mouse visual cortex. J. Neurosci. 31, 18506–18521 (2011).

23. Dipoppa, M. et al. Vision and locomotion shape the interactions between neuron types in mouse visual cortex. bioRxiv 058396 (2016).

24. Pachitariu, M., Stringer, C. & Harris, K. Robustness of spike deconvolution for calcium imaging of neural spiking. bioRxiv (2017).

25. Friedrich, J., Zhou, P. & Paninski, L. Fast online deconvolution of calcium imaging data. PLOS Comput. Biol. 13, e1005423 (2017).

26. Podgorski, K. & Haas, K. Fast non-negative temporal deconvolution for laser scanning microscopy. J. Biophotonics 6, 153–162 (2013).

27. Vogelstein, J. T. et al. Fast nonnegative deconvolution for spike train inference from population calcium imaging. J. Neurophysiol. 104, 3691–3704 (2010).

28. Friedrich, J. et al. Multi-scale approaches for high-speed imaging and analysis of large neural populations. bioRxiv 091132 (2016).

29. Glickfeld, L. L., Andermann, M. L., Bonin, V. & Reid, R. C. Cortico-cortical projections in mouse visual cortex are functionally target specific. Nat. Neurosci. 16, 219–226 (2013).

30. Harris, K. D., Quiroga, R. Q., Freeman, J. & Smith, S. L. Improving data quality in neuronal population recordings. Nat. Neurosci. 19, 1165–1174 (2016).

31. Spaen, Q., Hochbaum, D. S. & Asín-Achá, R. HNCcorr: A Novel Combinatorial Approach for Cell Identification in Calcium-Imaging Movies. ArXiv Prepr. ArXiv170301999 (2017).

32. Freeman, J. Open source tools for large-scale neuroscience. Curr. Opin. Neurobiol. 32, 156–163 (2015).

33. Reynolds, S., Onativia, J., Copeland, C. S., Schultz, S. R. & Dragotti, P. L. Spike detection using FRI methods and protein calcium sensors: performance analysis and comparisons. in Sampling Theory and Applications (SampTA), 2015 International Conference on 533–537 (IEEE, 2015).

34. Podgorski, K. & Ranganathan, G. Brain heating induced by near-infrared lasers during multiphoton microscopy. J. Neurophysiol. 116, 1012–1023 (2016).

35. Aharon, M., Elad, M. & Bruckstein, A. K-SVD: An algorithm for designing overcomplete dictionaries for sparse representation. IEEE Trans. Signal Process. 54, 4311–4322 (2006).

